# Cognitive task information is transferred between brain regions via resting-state network topology

**DOI:** 10.1101/101782

**Authors:** Takuya Ito, Kaustubh R. Kulkarni, Douglas H. Schultz, Ravi D. Mill, Richard H. Chen, Levi I. Solomyak, Michael W. Cole

## Abstract

Resting-state network connectivity has been associated with a variety of cognitive abilities, yet it remains unclear how these connectivity properties might contribute to the neurocognitive computations underlying these abilities. We developed a new approach – information transfer mapping – to test the hypothesis that resting-state functional network topology describes the computational mappings between brain regions that carry cognitive task information. Here we report that the transfer of diverse, task-rule information in distributed brain regions can be predicted based on estimated activity flow through resting-state network connections. Further, we find that these task-rule information transfers are coordinated by global hub regions within cognitive control networks. Activity flow over resting-state connections thus provides a large-scale network mechanism for cognitive task information transfer and global information coordination in the human brain, demonstrating the cognitive relevance of resting-state network topology.

---

The human brain is thought to be a distributed information-processing device, its routes of information transfer constituting a core feature that determines its computational architecture. Many studies have used correlations among resting-state functional MRI (fMRI) time series to study functional connectivity (FC) in the human brain^1^ (see ref. 1 for review). It remains unclear, however, if these resting-state FC routes are related to the brain's routes of cognitive information transfer. Evidence that group and individual differences in resting-state FC correlate with cognitive differences^2–4^ suggests that there is a systematic relationship between resting-state FC and cognitive information processing. However, without linking FC to information transfer, it remains unclear whether or how resting-state FC might mechanistically contribute to neurocognitive computations. Additionally, while a number of studies have shown that task information representations are distributed throughout the brain^5–8^, such studies have yet to reveal how these distributed representations are coordinated, and how information in any one brain region is used by other brain regions to produce cognitive computations^9^. Other studies investigating interdependence of brain regions during tasks (rather than during rest) have typically emphasized statistical dependencies between regional time series^10–12^, rather than the mechanistic transfer of task-relevant information content (reflected in task activation patterns^13^) between those regions. Thus, it remains unclear whether or how the network topology described by either restingstate or task-evoked FC is relevant to the neurocognitive computations underlying task performance.

Here, we provide evidence for a network mechanism underlying the transfer and coordination of distributed cognitive information during performance of a variety of complex multi-rule tasks. Based on recent evidence that resting-state FC describes the routes of task-evoked activity flow^14^ (Fig. 1A) – the movement of task activations between brain regions – we hypothesized that resting-state network topology describes the mappings underlying task information (task-evoked activation pattern) transfer between brain regions. If true, this hypothesis implicates a network mechanism for an information-preserving mapping across brain regions involving communication channels^9,15^ described by resting-state network topology. Identifying such a mechanism would provide an important new window into the large-scale information processing architecture of the human brain.

**Figure 1.**
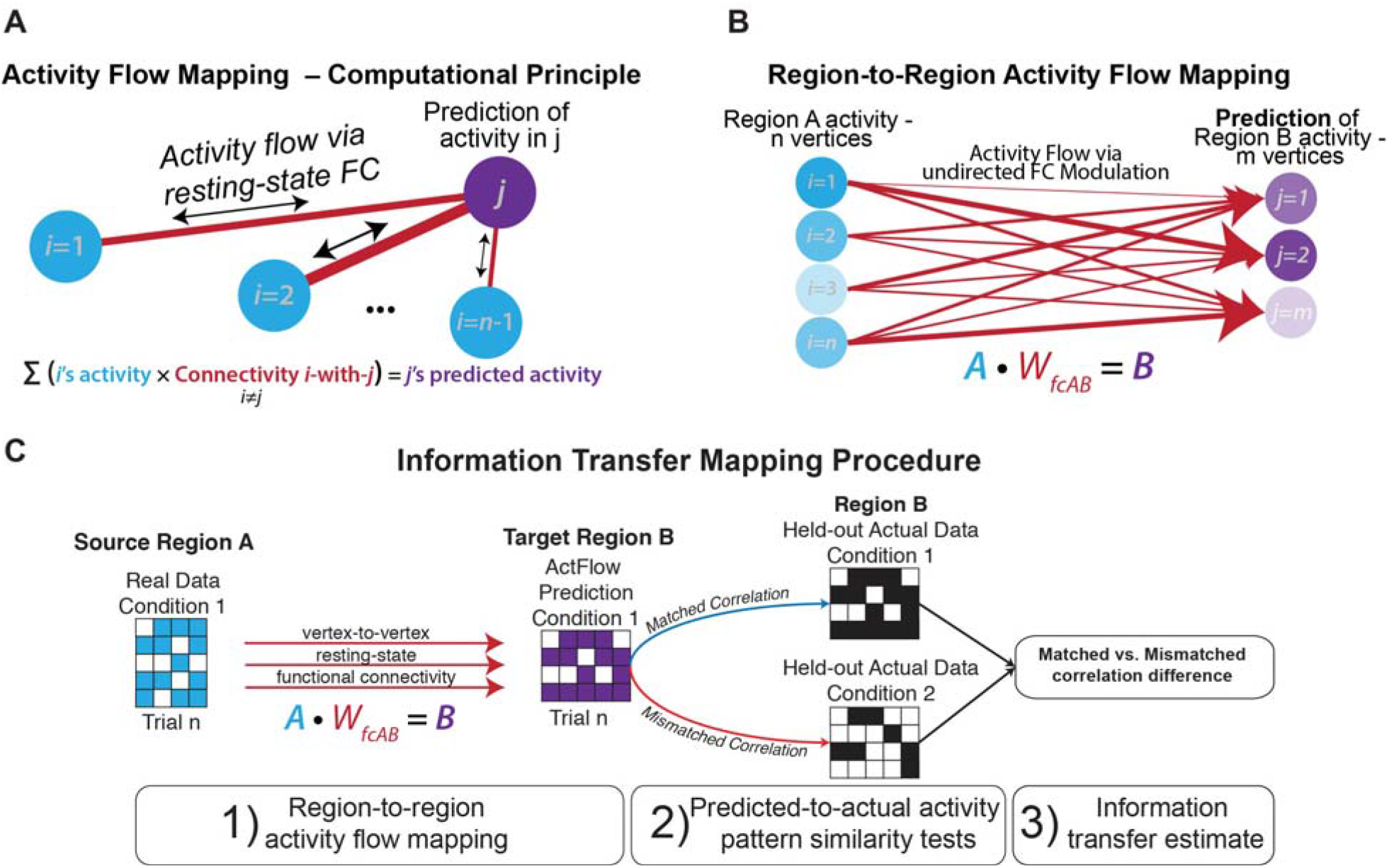
Measuring information transfer through activity flow mapping and cognitive task information decoding. **A)** Computational principle of activity flow mapping, as used by Cole et al. (2016)^14^. Adapted with permission from Cole et al. (2016). Activity in a held-out region is predicted by computing the linear weighted sum of all other regions' activity weighted by those regions' resting-state FC estimates with the held-out region. (The held-out region's activity is not included when computing the predicted activity of that region, thus avoiding a circular prediction.) **B)** Region-to-region activity flow mapping between vertices/voxels of isolated regions (“many-to-many” rather than “all-to-one” mapping of regions). Mathematically, we predict the activation pattern in Region B by computing the dot product of Region A's activation pattern vector with the vertex-to-vertex resting-state FC matrix between Region A and B. **C)** Information transfer mapping, which involves region-to-region activity flow mapping and representational similarity analysis (information decoding/classification) on held-out data. To test the transfer of task information from Region A to Region B, we compare the predicted activation pattern of Region B (mapped using Region A's activation pattern) to the actual task activation pattern of Region B for all task conditions using a spatial Spearman's rank correlation. For every prediction, spatial correlations to the task prototypes are computed and the information transfer estimate is measured by taking the difference of the correctly matched spatial correlation to the average of the incorrectly matched (mismatched) spatial correlations. Here we depict the approach for only two task conditions.

The current study focuses on fine-grained activation and FC topology, allowing us to infer the role of resting-state FC in carrying task-related information (represented by activation patterns^5–8^). This is, in turn, critical for testing a novel network mechanism in which resting-state FC topologies of cognitive control networks globally coordinate task-related information. Further, correspondence between resting-state FC topology and information-representing activation patterns would demonstrate the general mechanistic relevance of resting-state FC for information processing in the human brain.

Recent evidence suggests that resting-state FC reflects the human brain's invariant global routing architecture^16,17^. Supporting this, it has been demonstrated that most of the functional network topology variance present during task performance (80%) is already present during rest^18,19^. Thus, resting-state FC primarily reflects an intrinsic functional network architecture that is present regardless of cognitive context, given that there are only moderate changes to functional network organization across tasks^18,19^. We built upon these findings to test the hypothesis that intrinsic network topology describes the baseline network state upon which distributed cognitive information processing occurs.

Our hypothesis required an approach to empirically derive the mapping between information representations of pairs of brain regions, similar to identifying the transformation weights between layers in a neural network model^20^. The approach developed here contrasts with two previous approaches that describe the coordination of task-relevant information between brain regions. One of the previous approaches measures small shifts in task-evoked FC according to task-relevant content^10,12^. Another previous approach measures the correlation of moment-to-moment fluctuations in information content between regions^21^. Critically, these prior approaches primarily describe time-dependent statistical dependencies rather than suggest a large-scale mechanism by which task representations are mapped between brain regions. Thus, neither of these earlier approaches were appropriate for characterizing a network mechanism by which cognitive information is mapped between regions. Nonetheless, these past approaches were important for demonstrating the basic phenomenon of large-scale task information coordination, which we sought to better understand via the recently developed activity flow mapping approach^14^.

The hypothesis that fine-grained resting-state FC describes the representational mappings between brain regions during tasks is compatible with several recent findings. First, resting-state FC topology was recently shown to be highly structured and reproducible, forming clusters of networks consistent with known functional systems^22–24^. Second, as already mentioned, these resting-state networks are likely task-relevant given recent demonstrations that the network architecture estimated by resting-state FC is highly similar to FC architectures present during a variety of tasks^18,19^. Third, in addition to reflecting large-scale connectivity patterns, resting-state FC has been shown to reflect local topological mappings between retinotopic field maps in visual cortex, highlighting the specificity with which resting-state FC conserves functionally tuned connections^25,26^. Finally, resting-state FC has been shown to systematically relate to task-evoked activations, allowing prediction of an individual's task-evoked activations across a variety of tasks using that individual's resting-state FC^14,27^. This suggests a strong role for resting-state FC in shaping task activations – a core feature of our hypothesis that resting-state FC carries the fine-grained activation patterns that represent task-relevant information.

Traditional brain information mapping approaches localize task-related brain activity patterns. Because the experimenter is doing the information decoding, it is unclear whether (or how) that information is used for downstream processing by other brain regions. Thus, such approaches embody an experimenter-as-receiver framework, rather than a cortex-as-receiver framework, which estimates how brain regions send/receive information to/from other regions^9^. The proposed method – information transfer mapping – advances this perspective by analogizing resting-state connections with information channels. This allowed us to characterize whether distributed brain regions receive and decode task information from other brain regions via resting-state network connections, thus ascribing an information-theoretic description to resting-state network topology. Further, above-chance information transfers between two regions would indicate that the cognitive information in those brain regions is likely supported by the intrinsic network connectivity between them. Thus, information transfer mapping implicitly tests the cognitive relevance of resting-state FC topology.

Going beyond our general hypothesis, we additionally focus on the contribution of particular features of resting-state network topology in contributing to task-related information transfer. Recent studies have identified domain-general flexible hub networks that exhibit widespread resting-state FC and high activity during cognitive control tasks^10,28,29^. The strong involvement of these cognitive control networks – the frontoparietal network (FPN, which likely implements task sets^30^), cingulo-opercular network (CON, which likely implements task set maintenance^30^), and dorsal attention network (DAN, which likely implements top-down attentional processes^31^) – in cognitively-demanding processes suggests a role for flexibly transferring task information across regions and networks.

We sought to isolate cognitive representations that would likely involve cognitive control networks by using a cognitive paradigm that involves multiple features thought to be central to cognitive control. We used the Concrete Permuted Rule Operations (CPRO)^32^ paradigm (Fig. 2), which permutes rules in three different cognitive domains to produce dozens of unique task-sets. We predicted that cognitive control networks would flexibly represent task-rule information and transfer that information to other regions through their widespread intrinsic connections. The combination of experimental design and analytical framework allowed us to isolate cognitive operations and relate them to the neurobiological processes underlying activity flow mapping, thus targeting cognitive information transfer.

**Figure 2.**
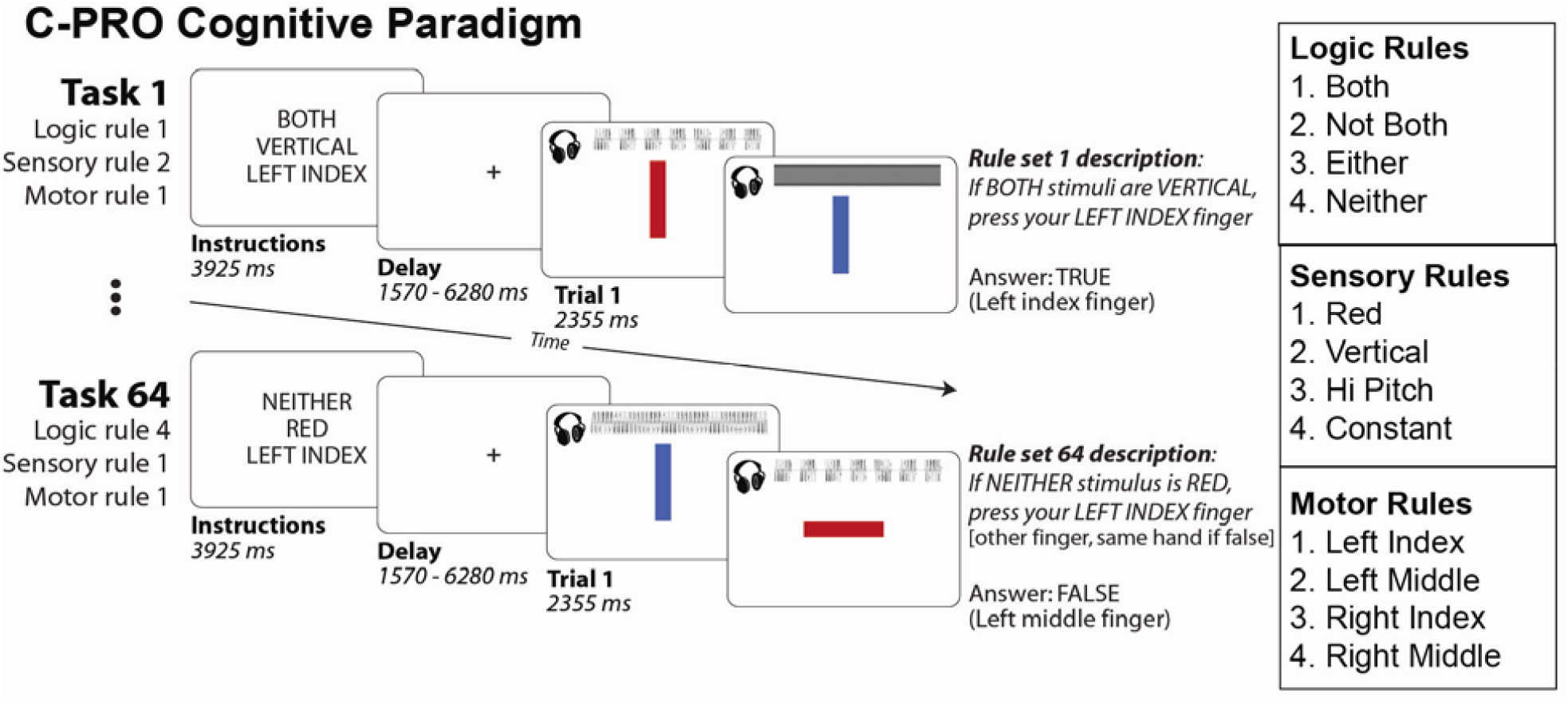
Concrete Permuted Rule Operations experimental paradigm. For a given task, subjects were presented with an instruction set (i.e., a task-rule set), in which they were presented with three rules each from a different rule domain (logic, sensory, and motor rule domains). Subjects were then asked to apply the presented rule set to two consecutively presented stimulus screens and respond accordingly. Auditory and visual stimuli were presented simultaneously for each stimulus screen. The auditory waveforms are depicted visually but were not presented visually to participants. A mini-block design was used, in which for a given set of instructions three trials were presented consecutively. The inter-trial interval was set to a constant 1570ms (2 TRs), with a jittered delay following the three trials prior to the subsequent task block (see Methods for more details). Task blocks lasted 28.26 seconds (36 TRs) each.

We began by replicating previously established properties of cognitive control networks, such as widespread resting-state FC^23,28^. We then used this replication to motivate a computational model that validates the effectiveness of the information transfer mapping procedure for estimating the role of resting-state network topology in transferring task information. Finally, we applied this framework to empirical fMRI data, allowing us to test our hypotheses that (1) resting-state FC describes channels of interregion/network task information transfer and (2) cognitive control networks play a role in transferring task information to other regions based on their intrinsic functional network properties. Our results show that the transfer of cognitive information could be reliably predicted using resting-state network topology, and cognitive control networks were especially involved in transferring information across multiple cognitive rule domains. Based on these results and a series of control analyses that confirmed that cognitive information transfer depends on precise resting-state network topology, we conclude that cognitive information used for task performance is transferred between brain regions via the functional network topology already present during resting state.

## Results

### Network organization of cognitive control networks

We began by establishing a strong basis for testing subsequent hypotheses regarding information transfer via cognitive control networks. Given the recent interest in reproducibility in neuroscience and other fields^33,34^, we replicated the hub-like characteristic of cognitive control networks^23,28,29,35^ before moving forward with analyses that build on these previous findings.

Using a recently developed set of functionally defined cortical regions^36^ (Fig. 3A), we tested whether cognitive control networks are global (connector) hubs. We quantified global hubs as having high between-network global connectivity (BGC) (see Methods) estimated during resting-state fMRI. We constrained our analyses to seven networks (Fig. 3A), identified by being replicated across multiple previously published functional network atlases^22–24^. We focused on BGC to reduce the bias toward larger mean connectivity (i.e., weighted degree centrality, or global brain connectivity^28^) for larger networks simply because they are larger^23,29^. We found that the top three networks with highest BGC estimated at rest were the three cognitive control networks: FPN, CON, and DAN (Fig. 3D; FPN greater than all non-cognitive control networks, with an averaged t_(31)_=9.52; CON greater than all non-cognitive control networks, with an averaged t_(31)_=12.33; DAN greater than all non-cognitive control networks, with an averaged t_(31)_=11.56; all family-wise error (FWE) corrected p < 0.0001). These results replicated previous results suggesting cognitive control networks are global hubs^23,28,29,35^, strengthening the basis for our hypothesis that cognitive control networks play a disproportionate role in shaping information transfer between regions throughout the brain. We test this hypothesis in a subsequent section, after establishing the validity of the newly-developed information transfer mapping procedure.

**Figure 3.**
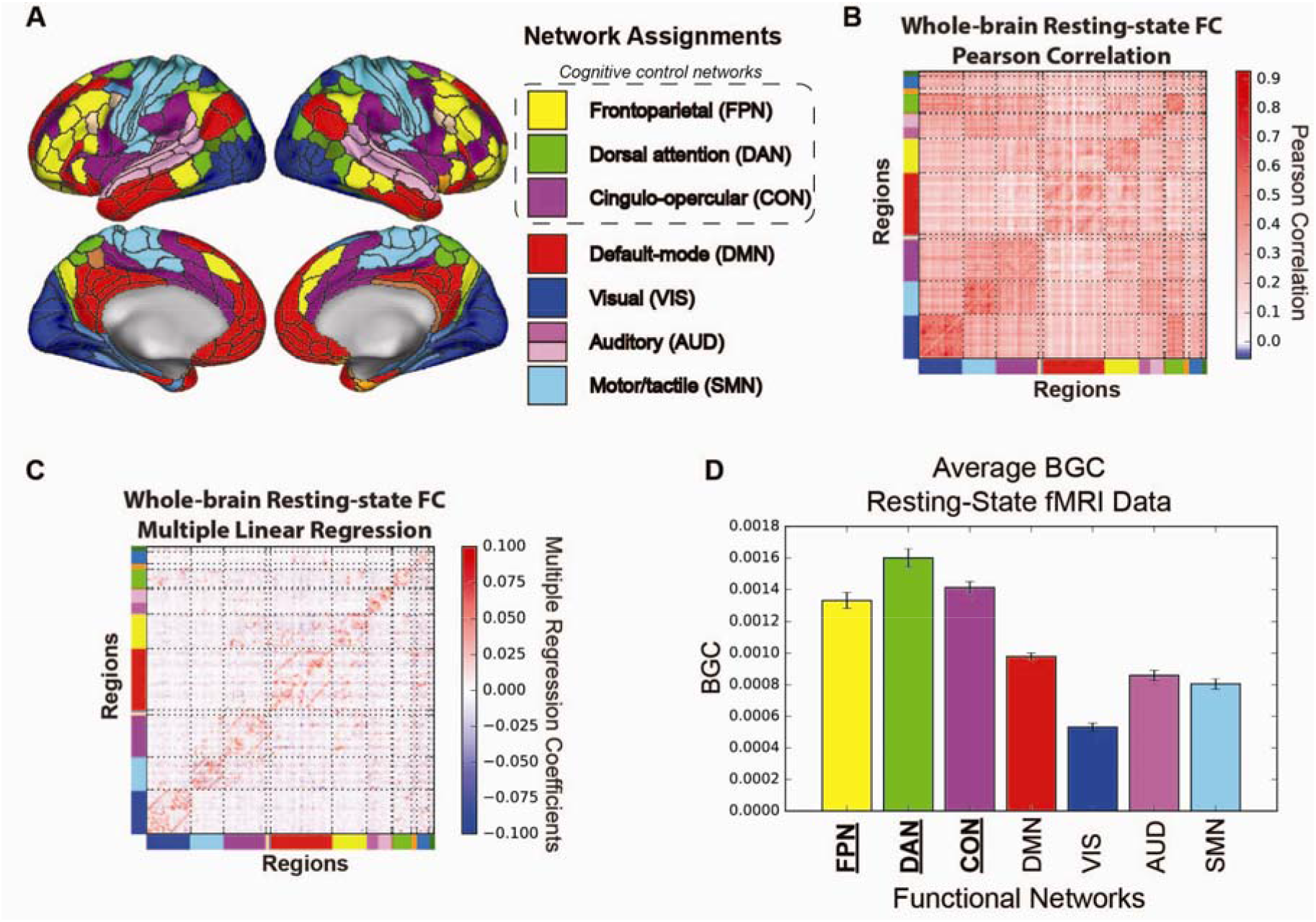
Large-scale network organization during rest. **A)** Using a recently released, multi-modal parcellation of the human cerebral cortex^36^, we assigned each region to a functional network using the Generalized Louvain method for community detection with resting-state fMRI data. We designated functional labels to seven networks that were replicated with other network assignments^22–24^. **B)** Whole-brain resting-state FC matrix computed using Pearson correlation between regions in panel A. Colors along the rows and columns denote network assignments from panel A. **C)** Whole-brain resting-state FC matrix computed using multiple linear regression. For every region's time series, we fitted a multiple linear regression model using the time series of all other regions as regressors of the target region. Multiple regression FC strongly reduced the chance that a connection was indirect, since FC estimates are based on unique shared variance. We used multiple regression FC for information transfer mapping, suggesting the estimated information transfers were likely direct rather than indirect. **D)** Averaged BGC of resting-state fMRI for each defined functional network. Cognitive control networks (underlined and in bold) had higher average BGC estimates relative to non-cognitive control networks (i.e., DMN and sensorimotor networks; FWE-corrected p<0.05). Error bars reflect across-subject standard error.

### Computational validation of information transfer mapping

We previously established that whole-brain activation patterns can be predicted based on activity flow over resting-state networks^14^. However, it remains unclear whether one region's cognitive information – coded as fine-grained activation patterns – can by predicted based on activity flow over resting-state FC. Such a demonstration would indicate that resting-state FC carries cognitive task information between brain regions (and networks). We tested this possibility by shifting from an “all-to-one” activity flow approach (i.e., predicting the activity level of a single brain region using the activity flow from all other brain regions; Fig. 1A) to modeling activity flow between a pair of regions (i.e., using the fine-grained activation pattern within one brain region to predict the fine-grained activation pattern within another region; Fig. 1B).

Testing our hypothesis required developing a new approach – information transfer mapping – which quantifies the amount of information transferred between pairs of brain regions over resting-state FC (Fig. 1B,C). Broadly, information transfer mapping tests the ability of resting-state FC topology (fine-grained connectivity patterns) to describe the mappings between cognitive-task-related activity patterns between pairs of brain regions. Specifically, each mapping (described by resting-state FC topology) must preserve the representational space between two regions, such that task-evoked information is decodable after the connectivity-based mapping. Beyond improving empirical understanding, this approach may have important theoretical implications given that it bridges biophysical (intrinsic FC) and computational (transformations between information-carrying activity patterns) properties into a convergent framework.

This approach (Fig. 1C) predicts the activation pattern in a target region based on a source region's activation pattern. This predicted activation pattern is then compared to the target region's actual activation pattern during the current task condition. The matched condition predicted-to-actual similarity is then compared to the mismatched condition predicted-to-actual similarity, with the difference in similarity quantifying the amount of task-specific information present in the prediction. Since the prediction was based on estimated activity flow over resting-state FC patterns, this allowed us to infer the amount of task-relevant information transferred via resting-state FC. Note that it was important to compare the predicted with the actual activation pattern in the target region to ensure that our prediction preserved the same representational geometry^37^ as the actual activation pattern.

We validated this approach using a simple abstract neural network model with one hub network and four non-hub networks (see Methods; Fig. 4A). This network organization was the basis for simulating fMRI dynamics during rest and task states, which allowed us to establish a “ground truth” to test the efficacy of the information transfer mapping procedure. This validation-via-modeling method was highly similar to the simple neural network model we previously used to validate the original activity flow mapping approach^14^. Using Wilson-Cowan type firing rate dynamics^38,39^, we simulated resting state and four distinct task states, simulated the transformation of the simulated neural signals to fMRI data (see Methods), and estimated resting-state FC (Fig. 4B) and task-evoked fMRI runs for each of the four task conditions (Fig. 4C). Note that we focused on network-to-network information transfer for our model validation (see schematic in Supplementary Fig. 1A), but later extended the approach to region-to region information transfer (see schematic in Fig. 6A).

**Figure 4.**
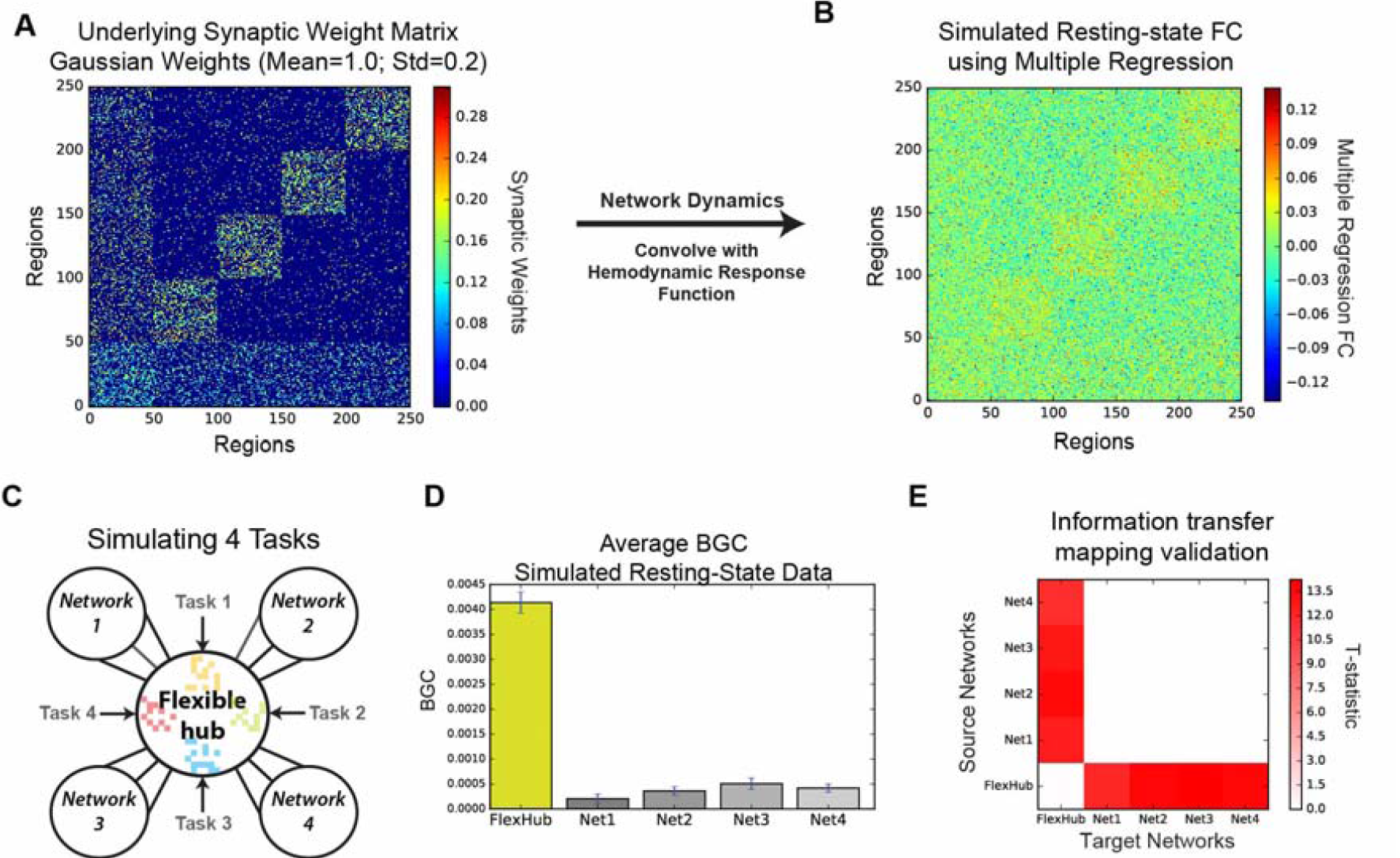
Computational validation of network-to-network information transfer mapping. **A)** Underlying synaptic weight matrix with four local networks and one hub network. We constructed an abstract neural network with a single hub network to see the relative effect of information transfer from the hub network to downstream local networks, similar to the hypothesized computational function of the cognitive control networks during task. **B)** Recovering large-scale synaptic organization via multiple regression FC estimates on a simulated resting-state time series. **C)** We simulated four ‘cognitive control tasks’ by stimulating four distinct ensembles of regions within the hub network. **D)** Increased BGC estimated at rest reflects underlying synaptic organization. Error bars represent across-subject standard error. **E)** Thresholded information transfer estimates between pairs of networks in a neural network model. Each row in the matrix corresponds to a source network from which we mapped activation patterns to other target networks using the information transfer mapping procedure (Fig. 1C). Each column in the matrix corresponds to a target network to which we compared the predicted-to-actual activation patterns. FWE-corrected thresholded T-statistic map with p < 0.05.

We found that simulated resting-state FC accurately reflected high BGC for the hub network (BGC statistically greater for the hub-network versus all other networks; averaged t_(29)_=21.14; FWE-corrected p<0.0001; Fig. 4D). Further, given the underlying synaptic connectivity structure (Fig. 4A) and the estimated intrinsic topology via restingstate FC (Fig. 4B,D), we hypothesized that information transfer to and from the hub network would reliably preserve task-specific information. Using the information transfer mapping approach (Fig. 1C; see Methods), we quantified the amount of information transfer via activity flow between every pair of networks (Fig. 4E). We found that information transfers to/from the flexible hub network and non-hub networks preserved task-specific representations (averaged information transfer estimate=0.13; averaged t_(29)_=11.86; FWE-corrected p<0.0001), while transfers between pairs of non-hub networks did not preserve statistically significant representations (averaged information transfer estimate=−0.0002; averaged t_(29)_=−0.02; averaged FWE-corrected p=0.91). We also found that these results were consistent with simulations where both top-down (hub network) and bottom-up (local network) stimulation occurred simultaneously (Supplementary Fig. 3; see Supplementary Methods). These results suggest that FC estimates obtained during simulated resting-state fMRI dynamics reflected underlying synaptic organization enough to describe the task-information-carrying mappings that govern activity flow between functional networks – a key assumption underlying our new approach.

These model simulations validated the plausibility of two hypotheses critical to the proposed information transfer mechanism: (1) Resting-state FC estimates characterize intrinsic FC (potentially reflecting aggregate synaptic connectivity) effectively enough to reflect underlying communication channel capacities; (2) Intrinsic FC describes the information-preserving mappings necessary to predict task-relevant activation patterns transferred from one region or network to another. Thus, these results validated the analytical basis of estimating information transfer via activity flow, which is applied to network-to-network and region-to-region information transfer mapping with empirical fMRI data below.

### Information transfer via resting-state network topology

We next applied the information transfer mapping procedure to real fMRI data, testing its ability to infer cognitive information transfer in the human brain. To test the hypothesis that cognitive control networks might widely distribute cognitive information via their resting-state network topology, we used an experimental paradigm with several features central to cognitive control to engage cognitive control networks. First, we used novel tasks given the need for control to specify behavior in such under-practiced scenarios^40,41^. Second, we used complex tasks given the need to deploy additional cognitive control resources when working memory is taxed^42^. Finally, we used a variety of abstract rules given that such rules are thought to be represented within cognitive control networks^5,43,44^. Using many fully-counterbalanced rules also allowed us to test our hypotheses across a variety of task conditions (while controlling for differences in sensory stimuli during trials). These features converged in the Concrete Permuted Rule Operations paradigm (C-PRO; Fig. 2). This paradigm was developed as part of this study, and is a modified version of the PRO paradigm^32^. We predicted that cognitive control networks would flexibly represent C-PRO rule information and transfer that information to other regions through their widespread intrinsic connections. For simplicity, we began with large-scale network-to-network information transfers. This involved quantifying information in large-scale functional networks based on patterns of region-level task activations (Supplementary Fig. 1; see Methods). In subsequent analyses we focused on region-to-region information transfers (based on patterns of voxel/vertex-level task activations).

As a prerequisite to running the network-to-network information transfer tests, we sought to first establish that task-rule information from the C-PRO paradigm (Fig. 2) was widely distributed across entire functional networks (Supplementary Fig. 1B). Logic rule information was significantly decodable in 6 out of 7 of the functional networks (averaged information estimate of significant effects=0.03; averaged significant t_(31)_=4.89; FWE-corrected p<0.01), with the somatomotor network (SMN) being the single network that did not contain decodable logic rule information (information estimate=0.007; t_(31)_=1.22; FWE-corrected p=0.58). Sensory rule information was significantly decodable in the FPN, DAN, CON, and visual network (VIS) (averaged information estimate=0.03; averaged t_(31)_=5.14; FWE-corrected p<0.001), and not decodable in the default mode network (DMN), auditory network (AUD), and SMN (averaged information estimate=0.003; averaged t_(31)_=0.83; averaged FWE-corrected p>0.11). Motor rule information was significantly decodable in the DAN, CON, and SMN (averaged information estimate=0.08; averaged t_(31)_=7.26; FWE-corrected p>0.0001), and not decodable in the FPN, DMN, VIS, and AUD (averaged information estimate=0.006; averaged t_(31)_=1.93; averaged FWE-corrected p>0.05). This allowed us to then evaluate whether significantly decodable representations of information were transferred to other functional networks.

In the logic rule domain, we identified information transfers between the FPN, CON, DMN, and AUD networks (Supplementary Fig. 1C; averaged information transfer estimate=0.009; averaged t_(31)_=4.73; FWE-corrected p<0.02). In the sensory rule domain, we found information transfers between the DAN and VIS in addition to the FPN, CON, and DMN (Supplementary Fig. 1D; averaged information transfer estimate=0.006; averaged t_(31)_=4.01; FWE-corrected p<0.05). Lastly, in the motor rule domain, information transfers were between the DAN, CON, and the SMN (Supplementary Fig. 1E; averaged information transfer estimates=0.011; averaged t_(31)_=5.37; FWE-corrected p<0.01). Further, to ensure that information transfers between pairs of networks was dependent on the precise network-to-network FC topology, we performed permutation testing, permuting FC patterns between pairs of networks (see Supplementary Methods). Indeed, after statistical testing, we found that information transfers were identical to our results with parametric statistical testing, suggesting that the observed information transfers were dependent on the specific resting-state network FC topology (Supplementary Fig. 4). These empirical network-to-network information transfers, along with their dependence on specific resting-state FC patterns, establish a role for resting-state network topology in transferring cognitive task information.

We next focused on region-to-region mappings that, unlike the network-to-network transfers, are based on fine-grained vertex-wise patterns (Fig. 6A). As a prerequisite to testing for information transfer between pairs of regions, we first needed to establish whether regions contained decodable task-rule representations. Thus, we first quantified the information content of each rule domain in the C-PRO paradigm (logic, sensory and motor rule domains) for each of the 360 regions using activation patterns (at the vertex level) with a cross-validated representational similarity analysis (see Methods). We found that logic rules were relatively distributed, with highest-quality representations in frontal and parietal cortices (averaged information estimate across significant effects=0.02; averaged t_(31)_=5.24; FWE-corrected p<0.05; Fig. 5A). Sensory rule information was also relatively distributed (averaged information estimate across significant effects=0.02; averaged t_(31)_=4.97; FWE-corrected p<0.05; Fig. 5B), though the highest-quality representations were predominantly in visual areas. Lastly, we found that motor rule representations were significantly more localized, with the highest-quality representations in the somatomotor network (averaged information estimate across significant effects=0.06; averaged t_(31)_=6.80; FWE-corrected p < 0.05; Fig. 5C). The existence of distributed task-rule information in multiple cortical regions allowed us to next assess how task-rule-specific information in one region might be transferred to other regions.

**Figure 5.**
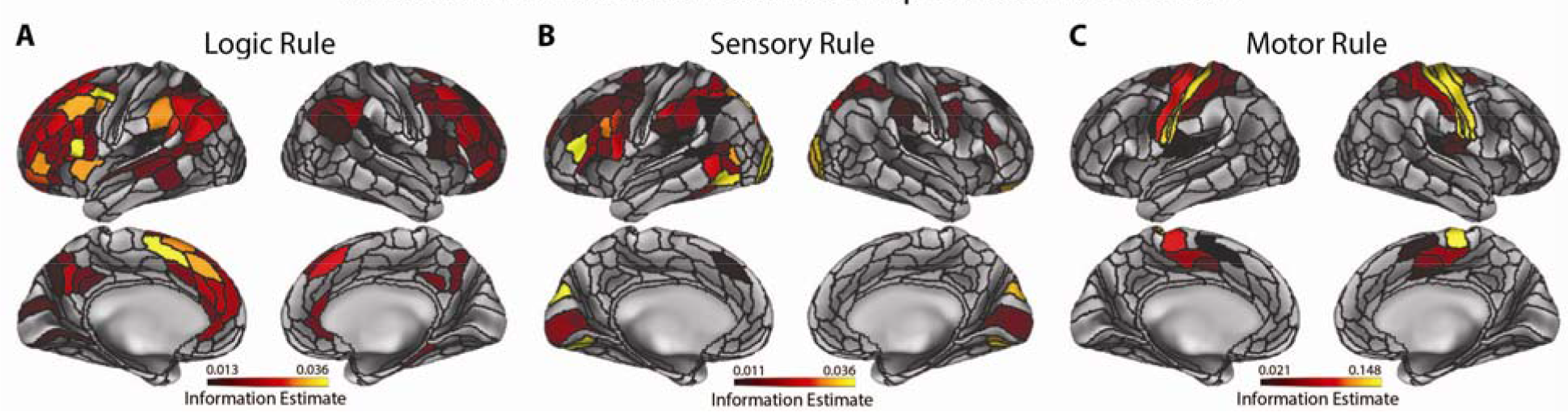
Information estimates of each region for each task-rule domain, prior to information transfer mapping. All reported results were statistically significant at FWE-corrected p<0.05. **A)** Thresholded whole-brain logic rule information estimate map. A cross-validated representational similarity analysis (quantifying degree of information representation; see Methods) for the logic rule domain was computed using vertices within every region. For each region, an average information estimate was computed for each subject, and a one-sided t-test was computed against zero across subjects. **B)** Thresholded whole-brain sensory rule information estimate map. As in the logic rule analysis, rule representations were highly distributed across the entire cortex, though representations were especially prominent in visual areas. **C)** Thresholded whole-brain motor rule information estimate map. Unlike the logic and sensory rule representations, motor rule representations were more localized to the motor/tactile network.

We next performed region-to-region information transfer mapping (Fig. 6). This approach utilized within-region vertex-level activation patterns along with vertex-to-vertex resting-state FC between regions to predict information content in each region (Fig. 6A; also see Methods). We performed this procedure for every pair of 360 regions, and visualized our results as a 360×360 matrix for each rule domain (Fig. 6B,D,F). However, given the difficulty in visually interpreting information transfers between every pair of regions (due to sparseness), we collapsed the region-to-region information transfer matrix by network to better visualize statistically significant region-to-region information transfers at the network level (Fig. 6C,D,G; see Supplementary Fig. 2A-C for all 14 networks). In addition, to see the relative anatomical position of regions that transferred information (i.e., source regions), we computed the percent of statistically significant transfers from each cortical region for each rule domain, and plotted these percentages on the cortical surface (Fig. 7A-C).

**Figure 6.**
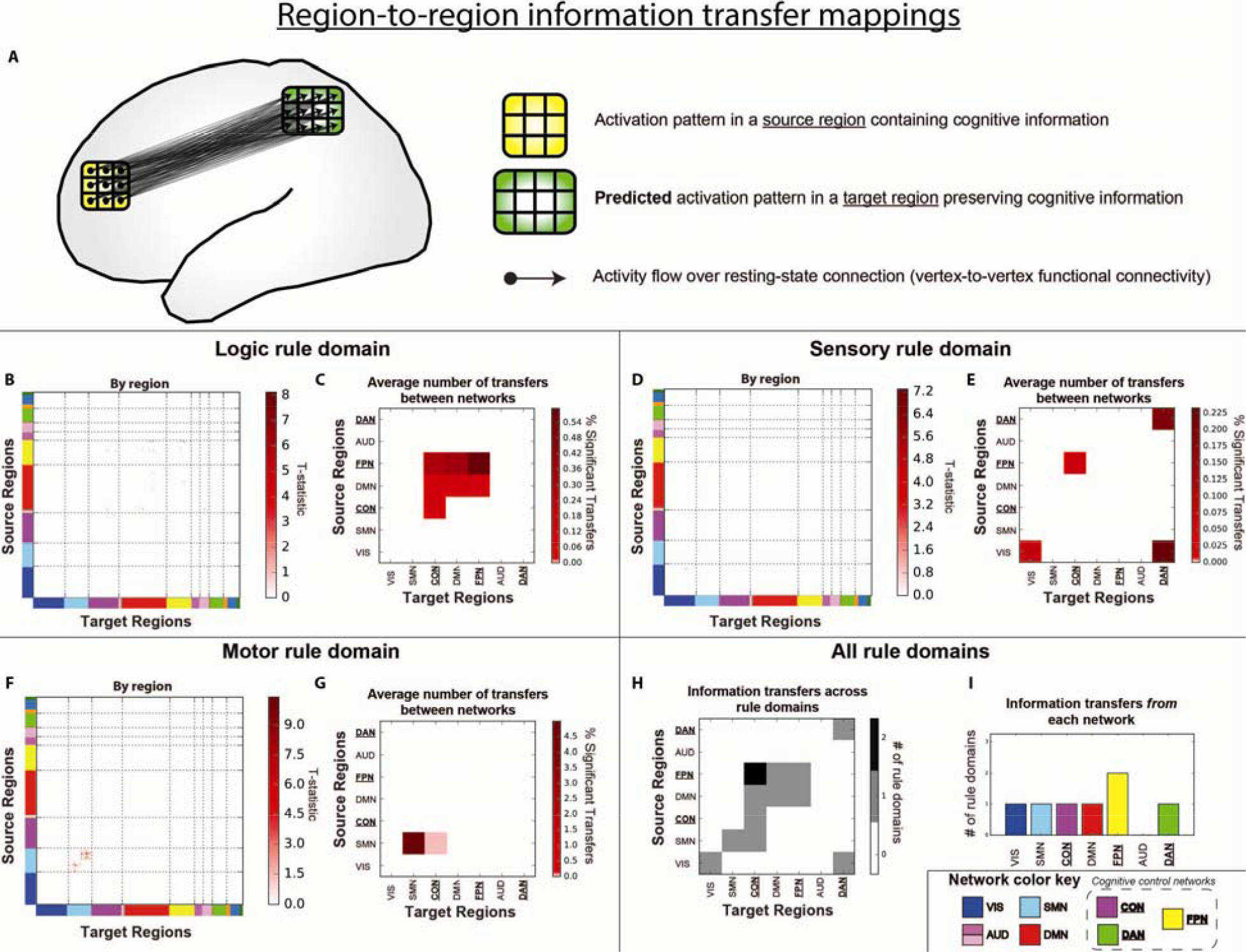
Information transfer mappings between all pairs of regions. All reported results were statistically significant at p<0.05 (FWE-corrected). (See Supplementary Fig. 6 for results with an FDR-corrected p<0.05 threshold.) **A)** Region-to-region information transfer mapping used the vertex-level activation pattern within one brain region and the fine-grained region-to-region resting-state FC topology to predict the vertex-level activation pattern in another brain region. **B)** Logic rule region-to-region information transfer mapping. **C)** Average number of statistically significant region-to-region transfers by network affiliations. To better visualize and assess how region-to-region transfer mappings may have been influenced by underlying network organization, we computed the percent of statistically significant rule transfers for every network-to-network configuration (i.e., the percentage of region-to-region transfers from a network A to a network B). (Note that visualizations for the full 14 network partition can be found in Supplementary Fig. 2.) Cognitive control networks are underlined. Information transfer of logic rule information was distributed across frontal and parietal cortices. **D,E)** Statistically significant sensory rule region-to-region information transfers. Region-to-region information transfers were substantially sparser for sensory rule mappings, but involved DAN and VIS regions. **F,G)** Statistically significant motor rule region-to-region information transfers. Motor rule mappings were noticeably more localized within the motor network. **H)** Statistically significant information transfers between regions grouped by network affiliation across rule domains. Across the three rule domains (panels C, E, and G) we counted the number of rule domains information was transferred between networks. **I)** We performed a similar analysis as in panel H, but counted the number of rule domains a network contained a region that transferred information (as a source region) across the three rule domains.

Overall, region-to-region information transfers were detected (FWE-corrected p<0.05) for all three task rule domains, as described in detail below. However, given the conservative nature of FWE correction, we also provide region-to-region information transfer results for false discovery rate^45^ (FDR) corrected p<0.05 thresholds, which potentially reduced false negatives but increased false positives (Supplementary Fig. 6 & 7). We found that with FDR correction, information transfers between regions were significantly more distributed (particularly in the logic rule domain). In both cases, these findings support the hypothesis that resting-state FC topology describes the channels of information transfer across multiple functional networks and across multiple task-content domains.

For logic rule mappings, while information transfers were highly distributed, most statistically significant region-to-region information transfers predominantly involved the FPN and other frontoparietal regions (averaged information transfer estimate across significant effects=0.02; averaged t_(31)_=6.26; FWE-corrected p<0.05). In particular, regions within the FPN transferred information to other regions in the FPN, as well as regions in other domain-general networks (CON and DMN) (Fig. 6C). Further, source regions involved in the transfer of logic rule information were left-lateralized for FWE corrected p<0.05 (Fig. 7A), although FDR-corrected p<0.05 thresholds showed more distributed source regions across bilateral frontal and parietal cortices (Supplementary Fig. 7A). In both cases, these findings suggest that the FPN uses intrinsic FC topology to distribute abstract (e.g., logic) rule information broadly for task set implementation and maintenance.

**Figure 7.**
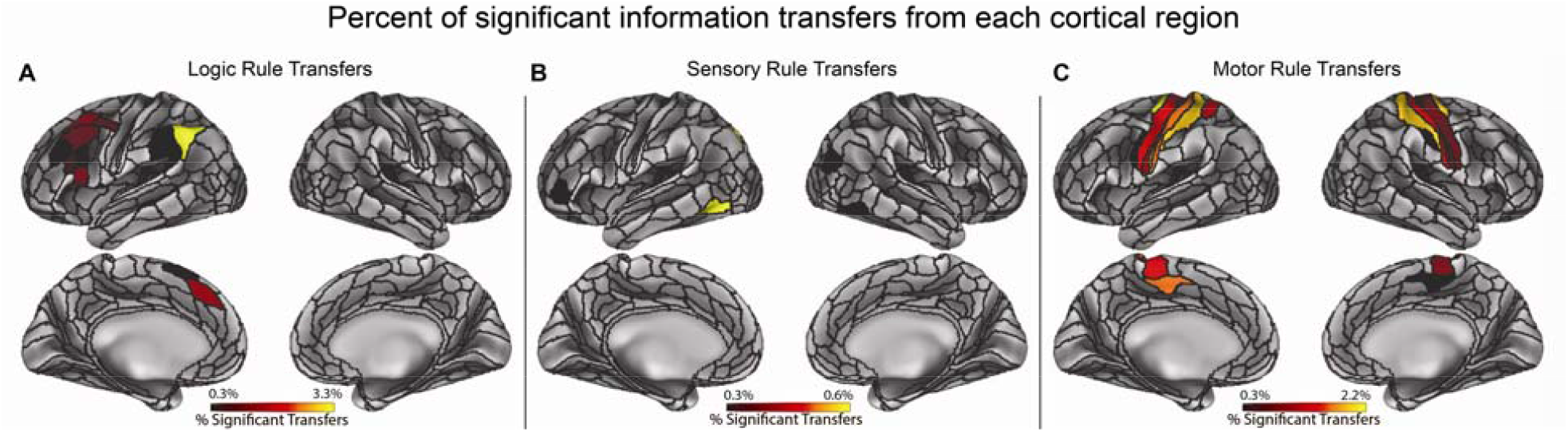
Percent of statistically significant information transfers from each cortical region. All reported information transfers were statistically significant at p<0.05 (FWE-corrected). (See Supplementary Fig. 7 for results with an FDR-corrected p<0.05 threshold.) **A)** Percent of statistically significant information transfers from each region for the logic rule domain. Percentages were computed by taking the number of significant transfers from each region, and dividing it by the total number of possible transfers from that region (359 other regions). Information transfers were relatively distributed, yet were predominantly from frontal parietal cortices. **B)** Percent of statistically significant information transfers from each region for the sensory rule domain. Information transfers were much sparser than in the logic rule domain. Most transfers were from higher-order visual areas and the DAN. **C)** Percent of statistically significant information transfers from each region for the motor rule domain. Transfers were predominantly from the motor network.

For sensory rule mappings, we found high specificity and sparseness of region-to-region task information transfers (averaged information transfer estimate across significant effects=0.01; averaged t_(31)_=6.15; FWE-corrected p<0.05; Fig. 7B). Most notably, we found that sensory rule representations are predominantly transferred within and between the DAN and VIS networks, as well as the FPN and CON (Fig. 6D,E). Previous studies have implicated a prominent role of the DAN and VIS in attentional processing of sensory information, consistent with the observed information transfers^31^. These findings suggest sensory rule information may be transferred between cognitive control networks, with transfers between regions in the DAN and VIS implementing these top-down information transfers.

Lastly, we found the most information transfer specificity for motor rule information (averaged information transfer estimate across significant effects=0.09; averaged t_(31)_=7.38; FWE-corrected p<0.05), consistent with the relatively localized representations of motor rule information (Fig. 5C). In particular, transfer of motor rule information largely involved transfers from regions in the SMN (Fig. 7C), while between-network information transfer with the SMN primarily involved the CON (Fig. 6G).

We next characterized the rule-domain generality of information transfers between specific networks. We found that regions within the FPN transferred rule information to the CON across two out of the three rule domains (Fig. 6H; see Supplementary Fig. 2D for all 14 networks). In addition, using an FDR-corrected threshold of p<0.05, we found statistically significant information transfers from FPN to CON for all three rule domains (Supplementary Fig. 6D). This is consistent with theories suggesting the FPN coordinates with CON to maintain and implement task sets^30^.

We next tested for networks that consistently transferred information across all rule domains, regardless of the target region's network affiliation. We found that regions in the FPN were consistently involved in transferring information to other regions in two rule domains (Fig. 6H). When using FDR to correct for multiple comparisons, we found that the FPN, DAN, and DMN transferred task information in all three rule domains (Supplementary Fig. 6E). We next assessed whether a single region transferred information across multiple rule domains. We found that no individual region consistently transferred task-rule information across the rule domains with either FWE or FDR correction, which suggests that unique sets of regions within each network were involved in transferring distinct types of cognitive information. This suggests that the regions within the FPN (and the DAN and DMN for FDR-corrected p<0.05 significance testing; Supplementary Fig. 6E) collectively act as flexible hub networks to communicate task-rules in different cognitive domains. Thus, the FPN likely plays an important role in task-rule transfers, regardless of cognitive domain.

These results uncover two key findings: (1) resting-state network topology describes the mappings likely underlying information transfer across distributed regions and functional networks, and (2) cognitive control networks likely play especially important roles in transferring a wide-range of task-rule information during complex cognitive tasks.

### Behavioral relevance of cognitive information transfer

We next tested whether estimated information transfers are predictive of task performance, demonstrating a likely role of information-pattern transfers in supporting task performance. Given that successful task performance required cognitive encoding of all three rule types (i.e., logic, sensory, and motor rules), we hypothesized that information transfer of all three rules were important to performing a task correctly. We therefore constructed a decoder using multiple logistic regression that was trained on the miniblock-to-miniblock information transfer estimates for all three rule types, and predicted the overall accuracy for held-out miniblocks (i.e., predicted a 1 if greater than 50% of trials were performed correctly within a miniblock, and 0 otherwise). Successful decoding of task performance using information transfer between pairs of regions would suggest that task performance depends in part on the successful transfer of task-rule information between those regions.

We first sought to ensure that task-rule information coded in the activity patterns used for information transfer mapping could predict behavioral performance, as a prerequisite to performing the information transfer mapping procedure. Given our findings that transfers between the FPN and CON were involved in two out of the three rule domains and that the FPN and CON are known to be involved in task-set maintenance^46^ we constrained our search to regions within those two networks. We found that rule information estimates (see Supplementary Methods) in a single FPN region in the lateral prefrontal cortex (LPFC) could significantly decode task performance (decoding accuracy=52.6%; t_(31)_=3.97; FWE-corrected p=0.02).

We then used this region as a source region and decoded task performance using information transfer estimates (across all rule domains) for transfers to every other region in the FPN and CON. We found that information transfer estimates from the LPFC region to an FPN region in the orbitofrontal cortex (OFC) could decode miniblock task performance significantly above chance (decoding accuracy=53.2%; t_(31)_=4.76; FWE-corrected p=0.003; Supplementary Fig. 5). This result demonstrates that the transfer of cognitive task-rule information between the LPFC and OFC was significantly correlated with task performance. However, while we account for the imbalance of correct and error trials in our decoding model, given that the behavioral data contains significantly fewer incorrect versus correct miniblocks, we interpret these results cautiously. (On average, 85% of miniblocks were performed correctly.) It will be important for future work to investigate the behavioral relevance of information transfers using a dataset that contains more error trials, allowing for a more robust model fit to behavior. Nonetheless, the combination of linking resting-state FC topology to information transfers across multiple brain systems and multiple cognitive task domains as well as trial-by-trial task performance strongly supports a role for resting-state FC topology in cognitive information transfer and task information processing.

## Discussion

Studies from neurophysiology, fMRI, and computational modeling emphasize the distributed nature of information processing in the brain^47–49^. However, fMRI studies often decode cognitive information from brain regions^6^ without considering how other brain regions might utilize that information^9^. In other words, current neuroscientific findings emphasize an experimenter-as-receiver framework (i.e., the experimenter decoding information in a brain region) rather than a cortex-as-receiver framework (i.e., brain regions decoding information transferred from other brain regions)^9^. The current emphasis on the experimenter-as-receiver framework clashes with the traditional understanding of information communication described by Shannon's Information Theory^15^, which provides a general theory of communication through the representation and transmission of information-bearing signals. Thus, understanding how cortical regions receive information from other regions bridges a crucial gap in understanding the nature of information processing in the brain. In light of recent findings relating resting-state fMRI to task-evoked cognitive activations^14,27^, we hypothesized that resting-state FC describes the channels over which information can be communicated between cortical regions. Results strongly supported this hypothesis, suggesting that resting-state network topology describes the large-scale architecture of information communication in the human brain and demonstrating the relevance of resting-state network connectivity to cognitive information processing.

We developed a novel procedure to quantify information transfer between brain regions. The procedure requires an information-preserving mapping between a source region and a target region. In the neural network modeling literature, analogous mappings are typically estimated through machine learning techniques to approximate synaptic weight transformations between layers of a neural network (e.g., an artificial neural network model using backpropagation)^20^. However, given that artificial neural networks are universal function estimators^50^ and would therefore fit any arbitrary mappings, we opted to take a more biologically principled approach that relied on FC estimation. Specifically, we used evidence that patterns of spontaneous activity can be used to successfully estimate the flow of task-related activity in both local and largescale brain networks^14,51,52^ to obtain biophysically plausible, data-driven mappings between brain regions using resting-state fMRI. Thus, information transfer mapping unifies both biophysical and computational mechanisms into a single information-theoretic framework.

We used a computational model to validate the plausibility of this account of large-scale information transfer, finding that despite the slow dynamics of the blood-oxygen level dependent signal, resting-state FC with simulated fMRI accurately reflects the large-scale channels of information transfer. We then used empirical fMRI data to show that resting-state FC describes information-preserving mappings in cortex at two levels of organization: brain regions and functional networks. In other words, the connectivity-based mappings estimated via resting-state FC between a source and a target region preserved task information content (in the same representational geometry^37,53^). Note that the organization of activity patterns was necessarily distinct between brain regions (given their distinct sizes and shapes), such that accurately predicting activation patterns in a target region based on activity in a source region reflected accurate spatial transformation of information-carrying activity patterns between those brain regions. These findings suggest that resting-state FC estimates likely reflect the actual large-scale mappings that are implemented in the brain during task information transfer.

We used multiple regression (rather than standard Pearson correlations) to estimate resting-state FC for information transfer mapping. This decision was based on recent evidence that activations are better predicted when using multiple-regression FC as compared to Pearson-correlation FC^14^. Importantly, multiple-regression FC strongly reduces the chance that estimated information transfers are indirect, since this method fits all regions/vertices simultaneously to identify unique shared variance between each pair of regions/vertices. Given that brain systems contain redundant neural signals^54^, however, multiple-regression FC estimates may be overly conservative. It will therefore be important for future research to validate appropriate regularization approaches to reduce the false negatives induced by multiple-regression FC. We expect that such a validated regularization approach would likely reveal that cognitive information transfers are even more widespread throughout the brain than reported here.

The evidence that fine-grained resting-state FC describes the information-preserving mappings between regions is important for advancing neuroscientific theory in a number of ways. First, the present results provide an empirically-validated theoretical account for how cognitive representations in different regions are likely mechanistically related to one another. Second, these results confirm the base assumption that decodable representations in a brain region are utilized by other regions through a biologically-plausible construct – information transfer via fine-grained patterns of activity flow. Third, these results expand the functional relevance of decades of resting-state FC findings^1,55^, given that we demonstrated the ability to use resting-state FC to describe cognitively-meaningful fine-grained relationships between brain regions. Importantly, our modeling and empirical results showed that the topological organization of the intrinsic connectivity architecture described inter-region information-preserving mappings. Further supporting this conclusion, we verified via permutation testing that fine-grained FC topology (rather than, e.g., overall mean FC) was essential for the observed information transfer results.

Previous studies have focused on the role of task-evoked FC in shifting distributed task representations^10,11^. We recently built on such findings to develop a flexible hub account of distributed task set reconfiguration via cognitive control networks^10,56^. The present results advance these findings by describing a network mechanism involving resting-state FC topology (and cognitive control network hubs) in transferring task representations throughout cortex. Importantly, recent findings have demonstrated that task-evoked FC changes tend to be small relative to resting-state FC topology^18,19^. This suggests that the resting-state FC topology investigated here likely carries the bulk of the task-relevant information transfers, with task-evoked FC alterations to this topology contributing only small (but likely important) changes to this process.

The information transfer mapping approach involves estimating linear information transfer. Critically, however, neural information processing is thought to often depend on nonlinear transformations^57^, such as face-selective neurons in the ventral visual stream responding to whole faces but not facial components (e.g., eyes and ears)^58,59^. The present findings represent an important step toward understanding the network mechanisms underlying information transformations between brain regions, setting the stage for future research to identify the role of resting-state FC in nonlinear information transformations. This would go beyond the information transfer processes investigated here to better identify the role of resting-state FC in neural computation (not just communication).

In summary, we combined information decoding of brain activity patterns with resting-state FC to demonstrate how fine-grained intrinsic connectivity patterns relate to cognitive information transfer. Further, by estimating information transfer throughout cortex we found evidence that cognitive control networks play important roles in global transfer of cognitive task information. We expect that these findings will spur new investigations into the nature of distributed information processing throughout the brain, providing a deeper understanding of these fine-grained information channels estimated at rest and their contribution to task-relevant information transfers.

## Methods

### Participants

35 human participants (17 females) were recruited from the Rutgers University-Newark community and neighboring communities. We excluded three subjects, leaving a total of 32 subjects for our analyses; two subjects were excluded due to exiting the scanner early, and one subject was excluded due to excessive movement. Excessive movement was defined as 3 standard deviations from the mean, in terms of framewise displacement^60^. All participants gave informed consent according to the protocol approved by the Rutgers University Institutional Review Board. The average age of the participants was 20, with an age range of 18 to 29.

### Behavioral paradigm

We used the Concrete Permuted Rule Operations (C-PRO) paradigm (Fig. 2), which is a modified version of the original PRO paradigm introduced in Cole et al., (2010)^32^. Briefly, the C-PRO cognitive paradigm permutes specific task rules from three different rule domains (logical decision, sensory semantic, and motor response) to generate dozens of novel and unique task sets. This creates a condition-rich dataset in the task configuration domain akin in some ways to movies and other condition-rich datasets used to investigate visual and auditory domains^48,61,62^. The primary modification of the C-PRO paradigm from the PRO paradigm was to use concrete, sensory (simultaneously presented visual and auditory) stimuli, as opposed to the abstract, linguistic stimuli in the original paradigm. Visual stimuli included either horizontal or vertical oriented bars with either blue or red coloring. Simultaneously presented auditory stimuli included continuous (constant) or non-continuous (nonconstant, i.e., “beeping”) tones presented at high (3000Hz) or low (300Hz) frequencies. Fig. 2 demonstrates two example task-rule sets for “Task 1” and “Task 64”. The paradigm was presented using E-Prime software version 2.0.10.353^63^.

Each rule domain (logic, sensory, and motor) consisted of four specific rules, while each task set was a combination of one rule from each rule domain (Fig. 2). A total of 64 unique task sets (4 logic rules × 4 sensory rules × 4 motor rules) were possible, and each unique task set was presented twice for a total of 128 task miniblocks. Identical task sets were not presented in consecutive blocks. Each task miniblock included three trials, each consisting of two sequentially presented instances of simultaneous audiovisual stimuli. A task block began with a 3925 ms instruction screen (5 TRs), followed by a jittered delay ranging from 1570 ms to 6280 ms (2 – 8 TRs; randomly selected). Following the jittered delay, three trials were presented for 2355 ms (3 TRs), each with an inter-trial interval of 1570 ms (2 TRs). A second jittered delay followed the third trial, lasting 7850 ms to 12560 ms (10-16 TRs; randomly selected). A task block lasted a total of 28260 ms (36 TRs). Subjects were trained on four of the 64 task-rule sets for 30 minutes prior to the fMRI session. The four practiced rule sets were selected such that all 12 rules were equally practiced. There were 16 such groups of four task sets possible, and the task sets chosen to be practiced were counterbalanced across subjects. Subjects' mean performance across all trials performed in the scanner was 85% (median=86%) with a standard deviation of 8% (min=66%; max=96%). All subjects performed statistically above chance (25%).

### fMRI Acquisition

Data were collected at the Rutgers University Brain Imaging Center (RUBIC). Whole-brain multiband echo-planar imaging (EPI) acquisitions were collected with a 32-channel head coil on a 3T Siemens Trio MRI scanner with TR=785 ms, TE=34.8 ms, flip angle=55°, Bandwidth 1924/Hz/Px, in-plane FoV read=208 mm, 72 slices, 2.0 mm isotropic voxels, with a multiband acceleration factor of 8. Whole-brain high-resolution T1-weighted and T2-weighted anatomical scans were also collected with 0.8 mm isotropic voxels. Spin echo field maps were collected in both the anterior to posterior direction and the posterior to anterior direction in accordance with the Human Connectome Project preprocessing pipeline^64^. A resting-state scan was collected for 14 minutes (1070 TRs), prior to the task scans. Eight task scans were subsequently collected, each spanning 7 minutes and 36 seconds (581 TRs). Each of the eight task runs (in addition to all other MRI data) were collected consecutively with short breaks in between (subjects did not leave the scanner).

### fMRI Preprocessing

Imaging data were minimally preprocessed using the publicly available Human Connectome Project minimal preprocessing pipeline version 3.5.0, which included anatomical reconstruction and segmentation, EPI reconstruction, segmentation, spatial normalization to standard template, intensity normalization, and motion correction^64^. All subsequent preprocessing steps and analyses were conducted on CIFTI 64k grayordinate standard space for vertex-wise analyses and parcellated time series for region-wise analyses using the Glasser et al. (2016)^36^ atlas (i.e., one time series for each of the 360 cortical regions). We performed nuisance regression on the minimally preprocessed resting-state data using 12 motion parameters (6 motion parameter estimates plus their derivatives) and ventricle and white matter time series (extracted volumetrically), along with the first derivatives of those time series.

Task time series for task activation analyses were preprocessed in an identical manner to resting-state data. Task time series were additionally processed as follows: A standard fMRI general linear model (GLM) was fit to task-evoked activity convolved with the SPM canonical hemodynamic response function and the same 16 nuisance regressors as above. Block-by-block activity beta estimates were used for representational similarity analyses and information transfer mapping analyses. Task activity GLMs were performed at both the region-wise level and vertex-wise level for subsequent network-to-network and region-to-region information transfer mapping, respectively.

### FC estimation

Given the success of FC estimation using multiple linear regression in our previous study^14^, we employed multiple linear regression to estimate FC. To estimate FC to a given node, we used standard linear regression to fit the time series of all other nodes as predictors (i.e., regressors) of the target nodes. Using ordinary least squares regression, we calculated whole-brain FC estimates by obtaining the regression coefficients from the equation

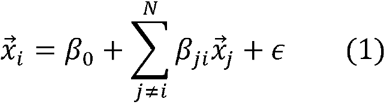

for all regions *x_i_* We define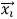 as the time series in region *x_i_*, *β_0_* as the y-intercept of the regression model, *β_ji_* as the F C coefficient for the *j*th regres regressor/region (which we use as row and the column in the FC adjacency matrix), and *∈* as the residual error of the regression model. *N* is the total number of regressors included in the model, which corresponds to the number of all other regions. This provided an estimate of the contribution of each source region in explaining unique variance in the target region's time series. This approach was used for region-to-region FC estimation, where the time series for each parcel was averaged across a given parcel's vertices prior to FC calculation. For this model *N* = 360, corresponding to the number of parcels in the Glasser et al. 2016 atlas^36^. Multiple linear regression FC is conceptually similar to partial correlation, but is actually semipartial correlation, as the estimates retain information about scaling a source time series (i.e., regressor time series) into the units of the to-be-predicted time series (i.e., predicted variable/target region).

For vertex-to-vertex FC estimation, due to computational intractability (i.e., more source vertices/regressors than time points), we used principal components regression with 500 principal components. This is the same form of regularized regression used in a previous study^14^ for voxel-to-voxel FC estimation. This approach involved reducing all source time series into 500 principal components and using the components as regressors to the target vertex. To reduce the possibility of spatial autocorrelation when estimating vertex-to-vertex FC, we excluded all vertices belonging to the same brain region/parcel as well as any vertices within 10mm of the border of that parcel in the principal components/regressors of the target vertex. (All vertices that fell within this criterion were given FC values of 0, preventing any vertices close to the target region from contaminating FC estimates.) Beta values obtained from the principal component regressors were then transformed back into the original 64k vertex space.

### Replication of network topological properties

We sought to replicate a key property of resting-state network topology using our novel network assignments of the Glasser et al. (2016) parcels – high global connectivity of cognitive control networks. We included only functional networks which coincided with the seven most replicable functional networks found in three previously published network atlases^12–14^: the frontoparietal network (FPN), the dorsal attention network (DAN), the cingulo-opercular network (CON), the default mode network (DMN), the visual network (VIS), the auditory network (AUD), and the somatomotor network (SMN). We measured the average between-network global connectivity (BGC) during resting-state FC, which was estimated using multiple linear regression (Fig. 3D). BGC connections were defined as all connections from the source region to target regions outside the source region's network. Mathematically, we defined each region's BGC as

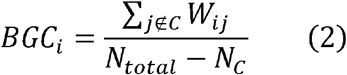

where *BGC_i_* corresponds to the BGC of region *i* in network *C,j ∉ C* corresponds to all regions not in network *C, W_ij_* corresponds to the FC estimate between regions *i* and *j*, *N_total_* corresponds to the total number of regions, and *N_c_* corresponds to the total number of regions in network *C* To compute the average BGC for a network *C* we averaged across all *BGC_i_* for *i ∊ C*

To statistically test whether the average BGC was different for a pair of networks, we performed a cross-subject paired t-test for every pair of networks. We corrected for multiple comparisons across pairs of networks using FWE permutation testing _65_.

### Neural network model

To validate our information transfer estimation approach we constructed a simple dynamical neural network model with similar network topological properties identified in our empirical fMRI data. We constructed a neural network with 250 regions, each of which were clustered into one of five network communities (50 regions per community). Regions within the same community had a 35% probability of connecting to another region (i.e., 35% connectivity density), and regions not assigned to the same community were assigned a connectivity probability of 5% (i.e., 5% out-of-network connectivity density). We selected one community to act as a “network hub”, and increased the out of-network connectivity density of those regions to 20% density. We then applied Gaussian weights on top of the underlying structural connectivity to simulate mean-field synaptic excitation between regions. These mean-field synaptic weights were set with a mean of 1.0/*√K* with a standard deviation of 0.2/*√K* where *K* is the number of synaptic inputs into a region such that synaptic input scales proportionally with the number of inputs. This approach was recently shown to be a plausible rule in real-world neural systems based on *in vitro* estimation of between-neuron synaptic-weight-setting rules^66^.

To simulate network-level firing rate dynamics, as similar to Stern et al. (2014), region *x_i_*'s dynamics for *i* = 1 … 250 obeyed the equation

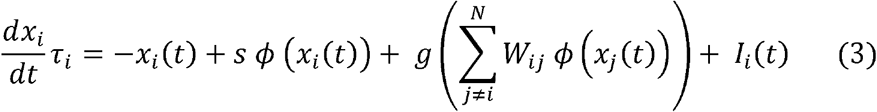

We define the transfer function ø as the hyperbolic tangent, *x_j_* the dynamics of region *j* = 1 … 250 for *i ≠ j, I_i_ (t)* the input function (e.g., external spontaneous activity alone or both spontaneous activity and task stimulation) for *i* = 1 … 250 *W* the underlying synaptic weight matrix, *s* the local coupling (i.e., recurrent) parameter *g* the global coupling parameter, and *τ_i_* the region's time constant. For simplicity, we set *s* = *g* = 1 and *τ_i_* = 10 ms, though we show in a previous study^14^ that the activity flow mapping breaks down for parameter regimes *s ≫ g*.

We first simulated spontaneous activity in our model by injecting Gaussian noise (parameter *I_i_(t)*; mean of 0.0, standard deviation 1.0). Numerical simulations were computed using a Runge-Kutta second order method with a time step of dt=10 ms. We ran our simulation for 600 seconds (10 minutes). To simulate resting-state fMRI, we then convolved our time series with the SPM canonical hemodynamic response function and down sampled to a 1 second TR, resulting in 600 time points. We then computed resting-state FC using multiple linear regression. To replicate the empirical data, we computed the BGC of the resting-state data (as in the empirical data; see equation 2) to validate that widespread out-of-network connectivity was preserved from synaptic to FC.

To model task-evoked activity, we simulated four distinct task conditions by injecting stimulation into four randomly selected but distinct sets of twelve regions in the hub network. Stimulation to the hub network was chosen to mimic four distinct top-down, cognitive control task rules. (See Supplemental Methods for further details.) We simulated 30 subjects worth of data, and generated figures using group t-tests and controlled for multiple comparisons using FWE-correction permutation tests^65^.

To perform network-to-network information transfer mapping in the model, we used the task-evoked activity (estimated by standard GLM beta estimates), and performed the information transfer mapping procedure between networks of regions using the resting-state FC matrix obtained via multiple linear regression. Network-to-network information transfer mapping is computationally identical to region-to-region information transfer mapping, and is described below.

### Computing information estimates for regions and networks

To compute the baseline (i.e., unrelated to FC) information content at the region level (Fig. 5), we performed a within-subject, cross-validated multivariate pattern analysis using representational similarity analysis for every Glasser et al. (2016) parcel (using the vertex-level multivariate activation pattern within each parcel). We estimated task-activation beta coefficients separately for each vertex within a region, and separately for each miniblock. Note that each miniblock was associated with a specific task-rule condition for each rule domain. Mathematically, we defined the information estimate of region B, as

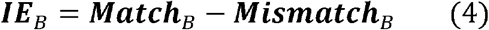

where **Match**_B_ and **Mismatch** correspond to the averaged Spearman rank correlation for matched and mismatched conditions, respectively. Specifically, we define **Match**_B_ and **Mismatch**_B_ as

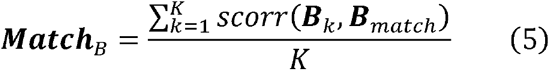

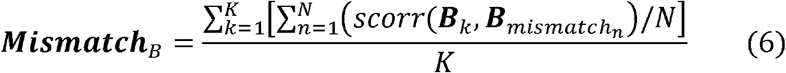

where *K* corresponds to the total number of miniblocks (in this paradigm, 128 miniblocks *K* corresponds to a Fisher *z*-transformed Spearman's rank correlation between two activation vectors, ***B**_k_* is the activation pattern in region B during block ***B**_match_* is the task-rule condition prototype (obtained by averaging across blocks of the same condition, holding out block *k*) of region B's activation pattern for which block *k*'s condition matches the condition prototype, and ***B**_mismatch_n__* as the task-rule condition prototypes for which block D's condition does not match. (In the present study *N = 3*, since each rule dimension has four task-rule conditions, and for a given miniblock there's one match and three mismatched conditions.) To avoid circularity, we performed a leave-four-out cross-validation scheme, holding out a miniblock of each task-rule. This ensured that miniblock ***B**_k_* was not included in constructing the condition prototype ***B**_match_* and that condition prototypes were each constructed using the same number of miniblocks. Prior to running the representational similarity analysis, all blocks were spatially demeaned to increase the likelihood that the representations we were identifying was a multivariate regional pattern (rather than a change in region-level mean activity). Use of Spearman's rank correlation also reduced the likelihood that the identified multivariate representation patterns were driven by mean activity changes or a small number of outlier values.

Statistical significance was assessed by taking a one-sided group t-test against 0 for each region's information estimate across subjects, since a greater than 0 difference of matches versus mismatches indicated significant representation of specific task-rules. All p-values were corrected for multiple comparisons across the 360 parcels using FWE-correction with permutation tests^65^, and significance was assessed using an FWE-corrected threshold of p<0.05.

(See Supplementary Methods for details on estimating network-level information estimates for Supplementary Fig. 1B.)

### Region-to-region information transfer mapping

We extended the original activity flow mapping procedure as defined in Cole et al. (2016)^14^ (Fig. 1A) to investigate transfer of task-related information between pairs of brain regions using vertex-wise activation patterns (i.e., regionto-region activity flow mapping; Fig. 1B). This involved predicting the activity of the vertices of a held-out target region based on the vertices within a source region. Mathematically, we define region-to-region activity flow mapping between regions A and B as

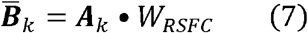

where ***B**_k_* corresponds to the predicted activation pattern vector for the target region B, ***A**_K_* corresponds to region A's activation pattern vector (i.e., the source region), *W_RSFC_* corresponds to the vertex-to-vertex resting-state FC between regions A and B, and the operator **•** refers to the dot product. This formulation allowed us to map activation patterns in one region's spatial dimension to the spatial dimension of another region.

To test the extent that task representations are preserved in the region-to-region multivariate predictions, we quantified how much information transfer occurred between the two regions. Briefly, information transfer mapping comprises three steps, illustrated in Fig. 1C: (1) Region-to-region (or network-to-network) activity flow mapping; (2) A cross-validated representational similarity analysis between predicted activation patterns and actual, held-out activation patterns; (3) Information classification/decoding by computing the difference between matched condition similarities and mismatched condition similarities. This final step produces an information transfer estimate. Mathematically, our information transfer estimate was derived using the exact formulation (equations 5 and 6) as our information estimate formula, but we substituted the target region's actual activation pattern ***B**_k_* for the target region's predicted activation pattern based ***B**_k_* on a connectivity-based transformation of source region A's activation pattern. (See Supplementary Methods materials for more details.)

Information transfer mapping was performed within subject between every pair of regions in the Glasser et al. (2016)^36^ atlas (360 regions in total). Statistical tests were performed using a group one-sided t-test (t > 0) for every pair-wise mapping. Our use of mismatched correlations as a baseline ensured that any positive information transfer estimates was a result of a task-rule-specific representation, rather than a task-general effect. Any information estimate that was not significantly greater than 0 indicated that the predicted-to-actual similarity was at chance (akin to chance decoding using classifiers). We tested for multiple comparisons using permutation testing^65^ for every region-to-region mapping, and significance was assessed using FWE-corrected pvalues with p < 0.05. Note that to avoid circularity for region-to-region information transfer mapping, any vertices in a source region that fell within a 10mm radius of the to-be-predicted target region (e.g., an adjacent region) would not contribute any activity flow to the to-be-predicted target region (see FC estimation Methods section for details). (See Supplementary Methods for further details.)

### Network-to-network information transfer mapping

Network-to-network information transfer mapping in both the computational model (Fig. 4E) and empirical data (Supplementary Fig. 1C-E) was performed in the same computational framework as above, though instead of predicting region-level activation patterns using vertex-level activation patterns, network-level activation patterns were predicted using region-level activations (averaging across vertices within a given region). (See Supplementary Methods for more details.)

### Behavioral relevance of information transfers

To characterize the behavioral relevance of information transfers, we performed a within-subject analysis to decode task performance using miniblock-by-miniblock information transfer estimates. We first sought to ensure that baseline miniblock information estimates could decode miniblock task performance within subjects prior to the information transfer mapping procedure. To perform a given task, knowledge of all three rule domains (i.e., logic, sensory, and motor rule domains) is required. Thus, we constructed a decoding model with logistic regression, training the model to decode the task performance of a given miniblock using the information estimates of a given brain region across all three rule domains. The model was tested using cross-validation in MATLAB using the glmfit function (with the logit link function). Miniblocks with over 50% of trials performed correctly were predicted as a 1, and 0 otherwise. However, to account for the imbalanced training data (on average, subjects performed 85% of trials correctly), we removed the intercept term β_0_ to center our predictions (as computed by a logistic function) at 0.5 (see Supplementary Methods for further details).

We applied our decoding model to all regions within the FPN and CON across subjects. For each region, we applied one-sided t-tests against chance (50%), and corrected for multiple comparisons using FWE-correction permutation tests^65^. We identified a single FPN region in the LPFC (left hemisphere region 80 in the Glasser et al. atlas; Supplementary Fig. 5) whose baseline information estimates predicted miniblock task performance.

We subsequently tested whether information transfer estimates from the LPFC region could predict task performance. We applied the decoding model to information transfer estimates across all rule domains for all information transfers from the LPFC region to all other FPN and CON regions. We performed one-sided t-tests against chance (50%) for each information transfer, and corrected for multiple comparisons using FWE-correction permutation tests^65^. We identified a single information transfer from the LPFC to the OFC (left hemisphere region 91; both FPN regions) that survived multiple comparisons with an FWE-corrected p<0.05. Surface visualizations for Supplementary Fig. 5 were made using Connectome Workbench software (version 1.2.3)^67^.

### Computational resources

Region-to-region information transfer mapping, vertex-to-vertex FC estimation, task-rule information estimation, and model simulations were performed on the Rutgers University-Newark supercomputer cluster (Newark Massive Memory Machine, NM3) using Python and MATLAB code.

### Data availability

We have included code demos with accompanying tutorial data for both our computational model and the empirical network-to-network information transfer mapping. We have also provided a GitHub repository with both MATLAB and Python code to run FWE-correction using permutation tests using the approach described in Nichols & Holmes, 2002^65^. Lastly, we have published all master scripts/jupyter notebooks used to generate results and figures in the manuscript. All other data presented in this study are available upon request.

Demo code for the information transfer mapping procedure is publicly available here: https://github.com/ColeLab/informationtransfermapping

Code for the FWE-correction via permutation testing is available here: https://github.com/ColeLab/MultipleComparisonsPermutationTstiing

## Acknowledgements

We thank Stephen J. Hanson, Catherine Hanson, and Gregg Ferencz in helping develop our MRI protocols for data collection. We also thank Merav Stern and Hiromichi Tsukada for helpful discussions regarding the computational model. We acknowledge support by the US National Institutes of Health under awards K99-R00 MH096801, R01 AG055556, and R01 MH109520. The content is solely the responsibility of the authors and does not necessarily represent the official views of any of the funding agencies.

## Author Contributions

T.I. and M.W.C. conceived of the study. T.I. performed data preprocessing, data analysis, and developed the computational model under the supervision of M.W.C. T.I., D.H.S., L.I.S., and M.W.C. designed the experimental paradigm, and M.W.C. programmed the experiment. K.R.K. and M.W.C. constructed network definitions. T.I., K.R.K., D.H.S., R.D.M., R.H.C., and L.I.S. performed the behavioral and fMRI experiments under the supervision of M.W.C. T.I. and M.W.C. wrote the manuscript.

